# SW#db: GPU-accelerated exact sequence similarity database search

**DOI:** 10.1101/013805

**Authors:** Matija Korpar, Martin Šošić, Dino Blažeka, Mile Šikić

## Abstract

The deluge of next-generation sequencing (NGS) data and expanding database poses higher requirements for protein similarity search. State-of-the-art tools such as BLAST are not fast enough to cope with these requirements. Because of that it is necessary to create new algorithms that will be faster while keeping similar sensitivity levels. The majority of protein similarity search methods are based on a seed-and-extend approach which uses standard dynamic programming algorithms in the extend phase. In this paper we present a SW#db tool and library for exact similarity search. Although its running times, as standalone tool, are comparable to running times of BLAST it is primarily designed for the extend phase where there are reduced number of candidates in the database. It uses both GPU and CPU parallelization and when we measured multiple queries on Swiss-prot and Uniref90 databases SW#db was 4 time faster than SSEARCH, 6-10 times faster than CUDASW++ and more than 20 times faster than SSW.

## Introduction

Searching for protein homologous has become a daily routine for the many biologists. Popular BLAST tools (PSI/DELTA/BLASTP) [1–3] produce search results for a single query in less than a second and many bioinformatical tools have come to depend upon the BLAST tool family to find matches in the database of sequences. However, protein sequence databases grow at an unprecedented pace and very often we would like to find homologous of not one, but hundreds, thousands or even more queries. With existing tools, the extensive time cost for such a search can hinder the research. BLAST family of tools, not being naturally parallelisable, is unable to utilize the development of new hardware focused on a low level parallelism (inter-core and many-core architectures).

While the exact dynamic programming algorithms such as Smith-Waterman [4] provide optimal and thus more sensitive results, they are generally slower than BLAST algorithms. The main reason for the BLAST acceleration is the heuristic culling of candidate sequences in the first step of database search. In the second step, BLAST family tools still use exact algorithms for computing the alignment between the query and the sequences from the reduced database. This step is usually the slower one. However, due to database reduction the total running time is much faster.

In this work we present our implementation of the database search sequence similarity method based on standard dynamic programming algorithms for sequence alignment tailored to utilize both CUDA GPUs and CPU SIMD instructions. SW#DB is an extension of our previous work on GPU-enabled exact pairwise alignments for long sequences (1). It supports multiple GPU cards. This implementation is additionally optimised for multiple queries rendering it significantly faster than both the state-of-the-art CUDASW++ [5–7] GPU enabled database search tool and SIMD-optimized (Single Instruction Multiple Data) tools such as SSEARCH [8] and SSW [9]. Also, depending on the databases and queries this tool can perform either slightly slower or faster than BLASTP.

Although the running times of SW#DB for searching the whole database are comparable with the ones of the BLAST family tools, our main intention is to provide an open source library for the alignment step of the database search tools. To provide the tools with maximum freedom and flexibility, we provide the functionality of using a custom subset of seed sequences from which to generate alignments, enabling researchers to experiment and tailor the results to their need. SW#DB is optimised for the simultaneous computing of alignments and due to this architecture, much faster than the implementations used in BLAST tools. The library could be used not only for protein database search based on the local protein alignment, but it could be also be used for the global or semi-global alignment of both protein and nucleotide queries on databases.

## Design and Implementation

In database alignment one or more sequences, also called queries, are aligned to the database of sequences. Deterministic database alignment using Smith-Waterman algorithm can be divided into three phases, scoring, filtering and reconstruction. In the scoring phase only the alignment score of every query-database sequence pair is obtained. In the filtering phase obtained scores are sorted by their *e-value*, calculated in the way found in BLAST [1], and filtered by the e-value threshold and the maximum number of desired alignments. In the reconstruction phase pairs of query-database sequences that passed the filtering are fully aligned.

Smith-Waterman algorithm has quadratic time complexity for both the scoring and the reconstruction phase, linear memory complexity for the scoring phase and quadratic memory complexity for the reconstruction phase. Because of the linear memory complexity scoring phase is much more suitable for CUDA implementation. In addition to CUDA enabled optimization, SW# uses single instruction multiple data (SIMD) architecture on both CPU and GPU processors and utilizes multi core CPU architectures. SW# database module is also divided into three submodules, scoring, filtering and reconstruction. Scoring and reconstruction submodules are time consuming and will be further explained.

SW# database module scoring submodule depends on five methods for scoring pairs of sequences, three utilizing CUDA architecture and two utilizing only the CPU architecture. Methods utilizing CUDA architecture are: single thread, single sequence method, single thread, multiple sequences method and multiple threads, single sequence method. Methods that run on CPU are: single thread, single sequence method and single thread multiple sequence method.

Each of the CUDA enabled methods is divided into two phases, database preparation and scoring phase. In the database preparation phase, database is converted to a specific format that enables the CUDA methods to maximally utilize the parallelization architecture, and copied to the graphics card. In the scoring phase, methods use the prebuilt database to score against the query. All of the CUDA methods also utilize query profile optimization [10] to lower the number of slow memory reads from the global memory. Number of CUDA threads will be denoted as T, and number of blocks as B.

CUDA single thread methods align the database in sequence blocks where each block contains B * T sequences sorted by length. Each sequence in the block is stored vertically in the memory. All sequences shorter than the maximum sequence length in the block are padded with the neutral element to avoid branching in the parallelized method. Memory consumption of this format of the database is at most two times the size of the input database. This method performs best on shorter database sequences that don’t deviate as much in their length. In single thread single sequence method, at a time B * T sequences are scored. To lower the number of slow memory operations, four sequence characters are read at a time. For every char four scores from the query profile are loaded at a time. To further reduce the number of reads, for each sequence character the query profile scores are read twice. With these optimizations, method solves 32 cells with only 9 memory read operations. Single thread, multiple sequence method works in the same way, except it utilizes CUDA SIMD operations to score four sequences at a time. Disadvantage of the SIMD method is that it supports scores up to 127, because the computation it done with chars. Another disadvantage of this method is that CUDA cards bellow architecture 3.5 do not support CUDA SIMD. However, we found this method very useful because, in general, most of the scores are fewer than 127.

Opposed to single thread CUDA method, multiple threads single sequence method performs best on long database sequences. It stores the database in its original format. Additional memory is used for the communication between threads which sums it up to nine times the memory needed for the input database. This method scores B sequences at a time with T threads working on each sequence. Threads are computing the score by solving the scoring matrix antidiagonal in parallel, using the wavefront method. Threads communicate through a horizontal bus. To lower the number of memory operations each thread solves four rows of the scoring matrix. Threads need to be synchronized after each thread solves one cell in each of four rows.

CPU single thread, multiple sequence method utilizes SWIMD library. SWIMD library is a CPU only Smith-Waterman database scoring library that utilizes SIMD and AVX architectures to solve up to sixteen sequences at a time. Library offers a method that calculates the scores up to 127 and is much faster that the unbounded ones. Disadvantage of this method is that it requires a minimum of SIMD 4.0 architecture which is not available on older CPUs.

All of the currently mentioned methods are used for scoring. SW# library contains CPU implementations of scoring and reconstructions with no architecture dependencies. These methods utilize Ukkonen's banded algorithm [11] to lower the time complexity. SW# library also contains CUDA accelerated reconstruction methods described in [12]. In addition for SIMD enabled CPU architectures reconstruction utilizes the SWIMD library.

SW# database aligning method begins with CUDA database initialization. Database initialization is time consuming process because copying large amount of data on the graphics card takes time. For that reason, initialization is done once and the prepared database can be used for unlimited number of queries. If the required space to store the database is smaller than available CUDA memory, database is split into smaller parts and is solved part by part. Scoring process begins by dividing sequences on short and long according to predefined threshold N. In parallel two methods are called, one for long sequences and one for short sequences. Long sequences are solved using SWIMD or standard SW# CPU implementation, depending which is available. Short sequences are solved in parallel, longer short sequences are solved the same way as long sequences, and shorter are solved with single thread, single or multiple sequences method, depending which is available. CUDA method begins with shorter sequences and CPU method begins with longer sequences and when the two methods overlap, the method is over. Using two parallel methods eliminates importance of the N parameter. When scoring of short sequences is finished, CPU method solving long sequences is stopped. Rest of the long sequences are solved with multiple threads, single sequence CUDA method. If CUDA SIMD is available the whole process is first done using SIMD methods which calculate scores up to 127. After the whole process is done, all sequences with score equal to 127 are scored again without using SIMD.

SW# database aligning software is available as library through a C API, as well as a standalone executable. Additionally standalone executable, which utilizes MPI for database alignment on multiple CUDA powered nodes, is provided in the package. Except with the mentioned Smith-Waterman algorithm, SW# provides alignment using Needleman-Wunsch [13] and semi-global alignment algorithms. SW# library is intended for external usage in database aligning heuristics, as it provides simple, flexible and powerful API. Main advantage of the library is in preparing the GPU database in advance for multiple scoring. In case of multi query database alignment, this lowers execution time significantly. Library also provides methods for database alignment with indexes, method of only aligning the selected sequences from the database. These methods can be very useful when used with heuristic database alignment solutions, since almost all of the heuristic solutions rely on Smith-Waterman algorithm at some point.

## Results

To systematically compare the performance of SW#DB with BLASTP, SSW, CUDASW++ (versions 2.0 and 3.1) and SSEARCH we used a list of proteins of various lengths (Table 1) and ASTRAL dataset [14] as queries and swissprot and Uniref90 as databases. Tests were performed on two configurations: single-GPU (Intel® Core™ i7-4770K CPU, 32 GB RAM, NVIDIA GeForce GTX 780, 256 GB SSD) and multi-GPU (Intel(R) Core(TM) i7-3770 CPU, 16 GB RAM, 2 * NVIDIA GeForce GTX 690, 256 GB SSD). Since NVIDIA GeForce GTX 690 has two GPUs, the multi-GPU configuration has 4 GPUs.

**Table 1.** The list of Uniprot IDs and lengths of proteins used in performance testing.

| Uniprot ID | Length (residues) |
| --- | --- |
| O74807 | 110 |
| P19930 | 195 |
| B8E1A7 | 299 |
| Q3ZAI3 | 390 |
| P18080 | 513 |
| O84416 | 607 |
| A9BIH4 | 727 |
| Q2LR26 | 804 |
| B4KLY7 | 980 |
| Q5R7Y0 | 1465 |
| Q700K0 | 5124 |
| P0C6V8 | 6733 |
| P0C6W9 | 7094 |
| O01761 | 8081 |
| Q6GGX3 | 10746 |
| Q9I7U4 | 18141 |
| Q8WXI7 | 22152 |
| Q3ASY8 | 36805 |

Although SW#DB is primarily tailored for multiple queries, we tested performances for single queries as well. The results in Fig 1 and Fig 2 show that while for shorter queries, up to 600 residues long, CPU based tools BLASTP and SSEARCH are faster, for longer queries GPU based tools are comparable to BLASTP and up to 4 times faster than SSEARCH. The slower running times for shorter queries are expected due to the latency in transferring database to GPU. In addition for shorter queries the parallelization is not as efficient as for longer ones. These figures do not include results achieved by SSW, because it was much slower than other tools. It was 3 to 15 times slower than the second slowest tool, SSEARCH.

**Fig. 1.**
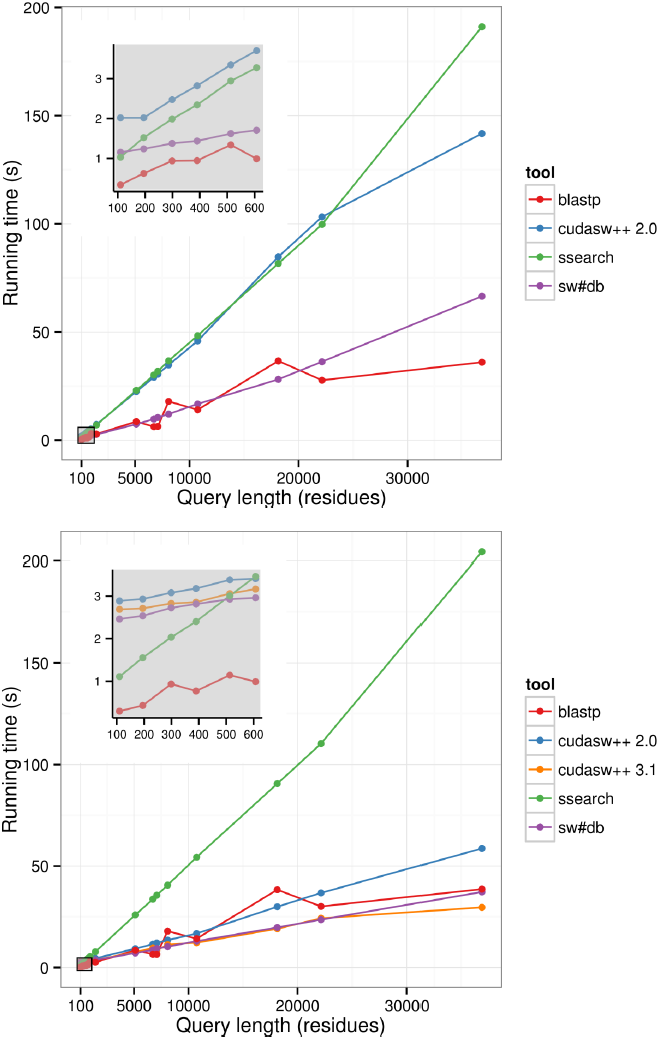
Comparison of SW#DB against BLASTP, CUDASW++ v. 2.0, CUDASW++ v. 3.1 and SSEARCH for queries of different length for Swiss-prot database. The insets show detailed results for shorter queries. Upper graph shows results for single-GPU machine (Nvidia GeForce GTX 780). Lower graph shows results for multiple-GPU machine (2 × GeForce GTX 690). We could not start CUDASW++ v.3.1 on single-GPU machine.

**Fig. 2.**
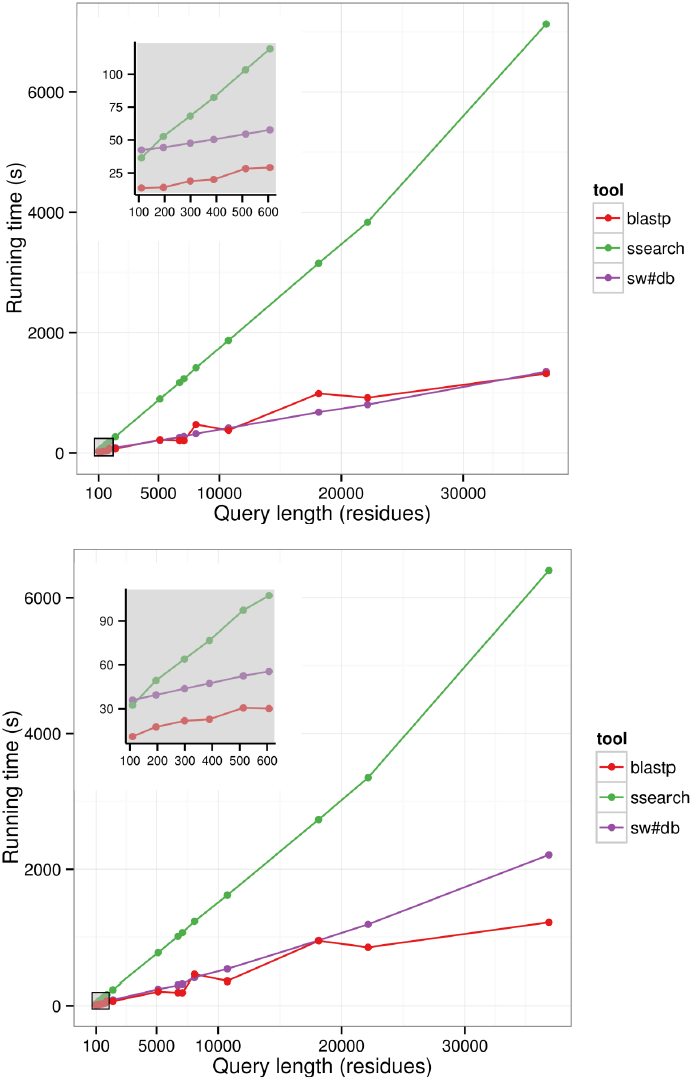
Comparison of SW#DB against BLASTP and SSEARCH for queries of different length for UniRef90 database. The insets show detailed results for shorter queries. Upper graph shows results for single-GPU machine (Nvidia GeForce GTX 780). Lower graph shows results for multiple-GPU machine (2 × GeForce GTX 690).

The real power of parallelization could be noticed for multiple queries. We used all programs to align ASTRAL database against UniprotKB/Swiss-prot and Uniref90 databases. The results are presented in Table 2. It shows that running times for BLASTP and SW#DB are comparable. For the smaller database (Swiss-prot) they are almost equal, while for the longer one (Uniref90) BLASTp is 1.7 times faster. In comparison with other parallelized exact algorithms SW#DB is 4 times faster than SSEARCH, 6-10 times faster than CUDASW++ and more than 20 times faster than SSW.

**Table 2.** Comparison of running times for SW#DB, BLASTP, CUDASW++ v2.0, CUDASW++ v3.1, SSW and SSEARCH using ASTRAL database as a query file. We could not run CUDASW++ 3.1 on the single-GPU machine (segmentation fault). Both versions of CUDASW++ could not run on Uniref90. We did not measure running time of SSW for Uniprot90 because it would last too long.

| Database | Configuration | Running time (s) |  |  |  |  |  |
| --- | --- | --- | --- | --- | --- | --- | --- |
|  |  | SW#DB | BLASTP | SSEARCH | CudaSW++ v2.0 | CudaSW++ v3.1 | SSW |
| Swiss-prot | Single-GPU<sup>1</sup> | 3523 | 3494 | 15123 | 23795 | - | 87118 |
| Uniref90 | Single-GPU<sup>1</sup> | 123581 | 73117 | 490543 | - | - | - |
| Swiss-prot | Multi-GPU<sup>2</sup> | 2709 | 3018 | 14088 | 32811 | 30545 | 85245 |

<sup>1</sup> Nvidia GeForce GTX 780 card
<sup>2</sup> 2×Nvidia GeForce GTX 690 cards

## Availability and Future Directions

The source code can be obtained from https://sourceforge.net/projects/swsharp/ and the tool is documented and rigorously tested. There are Windows and Linux releases. The further development of SW#DB will be focused on the better utilization of parallelization capabilities of both GPU and CPU and on the better load balancing between GPU and CPU.

## Conclusion

In this paper we present the SW#DB, a parallelised version of exact database search algorithms optimised for multiple queries. Although the emphasis is on the Smith-Waterman algorithm, other exact algorithms such as global and semi-global alignment are provided as well. SW#DB is parallelized on both GPU and CPU and it can run on multiple GPUs or in a cluster. The runningt times for large databases are comparable to times achieved by BLASTP and at least four times faster than state-of-the-art parallelized tools used for the same purposes such as SSEARCH, CUDASW++ and SSW. Although it could be used for the protein database search instead of BLASTP when the high sensitivity is required, our main intention was to build a library that could provide fast and exact alignment between queries and a reduced database for various bioinformaticstools.

## Acknowledgments

We would like to thank Ana Bulović for proofreading the manuscript.

